# *Contour*, a semi-automated segmentation and quantitation tool for cryo-soft-X-ray tomography

**DOI:** 10.1101/2021.12.03.470962

**Authors:** Kamal L Nahas, João Ferreira Fernandes, Colin Crump, Stephen Graham, Maria Harkiolaki

## Abstract

Cryo-soft-X-ray tomography is being increasingly used in biological research to study the morphology of cellular compartments and how they change in response to different stimuli, such as viral infections. Segmentation of these compartments is limited by time-consuming manual tools or machine learning algorithms that require extensive time and effort to train. Here we describe *Contour*, a new, easy-to-use, highly automated segmentation tool that enables accelerated segmentation of tomograms to delineate distinct cellular compartments. Using *Contour*, cellular structures can be segmented based on their projection intensity and geometrical width by applying a threshold range to the image and excluding noise smaller in width than the cellular compartments of interest. This method is less laborious and less prone to errors from human judgement than current tools that require features to be manually traced, and does not require training datasets as would machine-learning driven segmentation. We show that high-contrast compartments such as mitochondria, lipid droplets, and features at the cell surface can be easily segmented with this technique in the context of investigating herpes simplex virus 1 infection. *Contour* can extract geometric measurements from 3D segmented volumes, providing a new method to quantitate cryo-soft-X-ray tomography data. *Contour* can be freely downloaded at github.com/kamallouisnahas/Contour.

**Impact Statement:** More research groups are using cryo-soft-X-ray tomography as a correlative imaging tool to study the ultrastructure of cells and tissues but very few tomograms are segmented with existing segmentation programs. Segmentation is usually a prerequisite for measuring the geometry of features in tomograms but the time- and labour-intensive nature of current segmentation techniques means that such measurements are rarely across a large number of tomograms, as is required for robust statistical analysis. *Contour* has been designed to facilitate the automation of segmentation and, as a result, reduce manual effort and increase the number of tomograms that can be segmented. Because it requires minimal manual intervention, *Contour* is not as prone to human error as programs that require the users to trace the edges of cellular features. Geometry measurements of the segmented volumes can be calculated using this program, providing a new platform to quantitate cryoSXT data. *Contour* also supports quantitation of volumes imported from other segmentation programs. The generation of a large sample of segmented volumes with *Contour* that can be used as a representative training dataset for machine learning applications is a long-term aspiration of this technique.

## Introduction

The biology of cellular compartments has been extensively studied using high-resolution microscopy techniques. Transmission electron microscopy of thin sections of cells stained with heavy metals has been used for decades to produce images of intracellular ultrastructure and can resolve structures at the nanometer level^(1)^. For precise quantitation, cellular compartments of interest need to be delineated from the other ultrastructural features by segmentation. These features can be segmented manually by tracing the edges of features with Segmentation Editor in Fiji^(2)^, or with tools such as Amira (Thermo Scientific) that have ‘intelligent scissors’ that predict the boundaries of the object being traced by the user^(3)^. However, these manual processes are time-consuming and the boundaries of the segmented volumes are prone to human interpretation^(4)^. Automatic tools exist, but these also have limitations. For example, Bayesian matting, wherein a Bayesain framework is used to delineate foreground objects from the background based on pixel range, is less likely to successfully segment features with textured or thin edges^(5)^. Similarly, ‘magic wand’ segmentation, in which pixels of a given range of intensities are segmented if they are all connected, is less applicable to features with a broad range of intensities and where there is high noise in the background^(6,7)^. Watershed segmentation is often used to separate objects by estimating the boundaries between them based on the distances between their highest intensity maxima. However, the specificity of this technique is low in noisy datasets and can lead to over-segmentation, whereby many small segments are created within a single feature^(8,9)^. As a result, segmentation tools that use machine learning and deep neural networks to distinguish features of interest from the rest of the ultrastructure have been developed for electron microscopy (e.g. Unet, Ilastik)^(10–15)^. However, these tools require either a large representative training dataset or modified training for each micrograph.

The ultrastructural imaging technique known as cryo-soft-X-ray tomography (cryoSXT) has recently become accessible as a tool to cell biologists and pathologists to image the cellular compartments of unfixed whole cells in 3D^(16,17)^. Moreover, cryoSXT is being used as a correlative imaging technique with cryo-structured illumination microscopy (cryoSIM) to identify features in cellular ultrastructure^(18,19)^. X rays with a relatively low energy (∼0.5 keV)^(16)^, compared with those used for crystallography and medical imaging (∼5–30 keV)^(20,21)^, are used to illuminate the sample and transmission is reduced by absorption through carbon-rich structures, such as membranous cellular compartments. As a result, the signal in cryoSXT data appears dark due to X-ray absorption and the background appears light due to X-ray transmission. This technique is used to resolve cellular compartments to a theoretical resolution limit of 25 nm and produce 3D tomograms of whole-cell ultrastructure^(17)^. CryoSXT imaging of cells and tissues takes 5–20 minutes and thus a large set of tomograms—each containing cellular compartments that need to be delineated by segmentation—can be collected in a relatively short interval^(16)^. However, segmentation tools to mine information out of X-ray tomograms still need to be developed. One reason for this may be that X-ray tomograms are more difficult to segment than electron micrographs because the use of soft X rays to image the cell volume in 3D under near-native conditions produces higher noise and lower contrast than the heavy metal labelling used in electron microscopy^(22)^.

Although manual segmentation can be used to isolate features of interest, this is more time-consuming for 3D datasets that span the entire depth of the field of view within the cell^(4)^. The development of machine learning tools for cryoSXT data could increase the rate and efficiency of segmentation. However, the resolution, density and morphology of features can vary widely between cryoSXT datasets (e.g. depending on collection date, passage number of cultured cells, sample preparation strategy, etc.^(23)^), and this lack of consistency may complicate the use of machine learning tools to segment tomograms. Currently, there is a lack of training datasets for machine learning in the form of segmented volumes from multiple tomograms. SuRVoS has been developed to circumvent the need for training datasets in this form. Instead, individual frames are segmented and used to train segmentation of the whole tomogram^(4)^. However, this strategy involves training for each tomogram, which is time-consuming and does not keep pace with the high rate of cryoSXT tomogram acquisition.

Here we developed *Contour*, a semi-automated segmentation tool for cryoSXT. This tool can be used to segment high contrast features in cryoSXT tomograms, such as mitochondria, lipid droplets, and membranous features. This is achieved by a combination of thresholding based on the projection intensity (i.e. darkness) of the features and applying a width restriction based on the size of the features. This automated procedure can be performed globally (i.e. on the entire tomogram). Some features of interest may be excluded due to the strict width restriction, but segmentation of these features can be refined locally in smaller regions of interest. *Contour* was developed using Python 3.7 and is available for download on Github with example datasets included (github.com/kamallouisnahas/Contour). The segmentation approach used in *Contour* is faster than manual segmentation tools as it does not require laborious freehand drawing and interpolation like the Segmentation Editor available in Fiji^(2)^.

Extracting quantitative data from cryoSXT datasets is a current challenge and *Contour* can be used to measure the volume of segmented elements as well as their width along their longest axis. *Contour* was designed to be used alongside existing segmentation tools: for features that are difficult to segment based on projection intensity and width in *Contour* (e.g. cytoplasmic vesicles) other segmentation tools can be used to generate segmented volumes that can be imported into *Contour* for quantitation. We have used *Contour* in a recent preprint to study how the morphology of mitochondria and cytoplasmic vesicles change during infection with herpes simplex virus-1 (HSV-1)^(24)^. We generated multiple segmented volumes with *Contour* and found that mitochondria became more elongated and vesicles reduced in width as the infection progressed^(24)^. In this paper we discuss the algorithm and applications of this segmentation tool to cryoSXT data.

## Results

### The width of cellular compartments and the projection intensity of their voxels can be exploited for semi-automated segmentation

High-contrast cellular compartments in tomograms can be segmented by applying a threshold on voxel intensity. CryoSXT Z stacks were generated for segmentation using IMOD version 5.1.2^(25)^ with a back projection strategy, radial filtering in the form of a simultaneous iterations reconstruction technique (SIRT)-like filter being subsequently applied to reduce noise. Twenty iterations of the SIRT-like filter were applied to limit blurring and signal loss^(26)^. Mitochondria have a low voxel intensity (high X ray absorbance) compared with the cytosol and an arbitrary threshold range determined by trial and error was used to segment them in a U2OS cell from an 8-bit reconstructed tomogram (Fig. 1A)^(18)^. However, segmentation based solely on projection intensity was observed to be highly sensitive to voxel noise and non-specific features, such as the outline of the lipid droplets. In order to increase specificity, an additional segmentation parameter in *Contour* was used based on the width of the cellular compartments of interest (Fig. 1B). Segmentation was first performed on a complete reconstructed cryoSXT Z stack using the global segmentation algorithm in *Contour*. The segmentation was later refined in smaller regions using the local segmentation algorithm.

**Figure 1.**
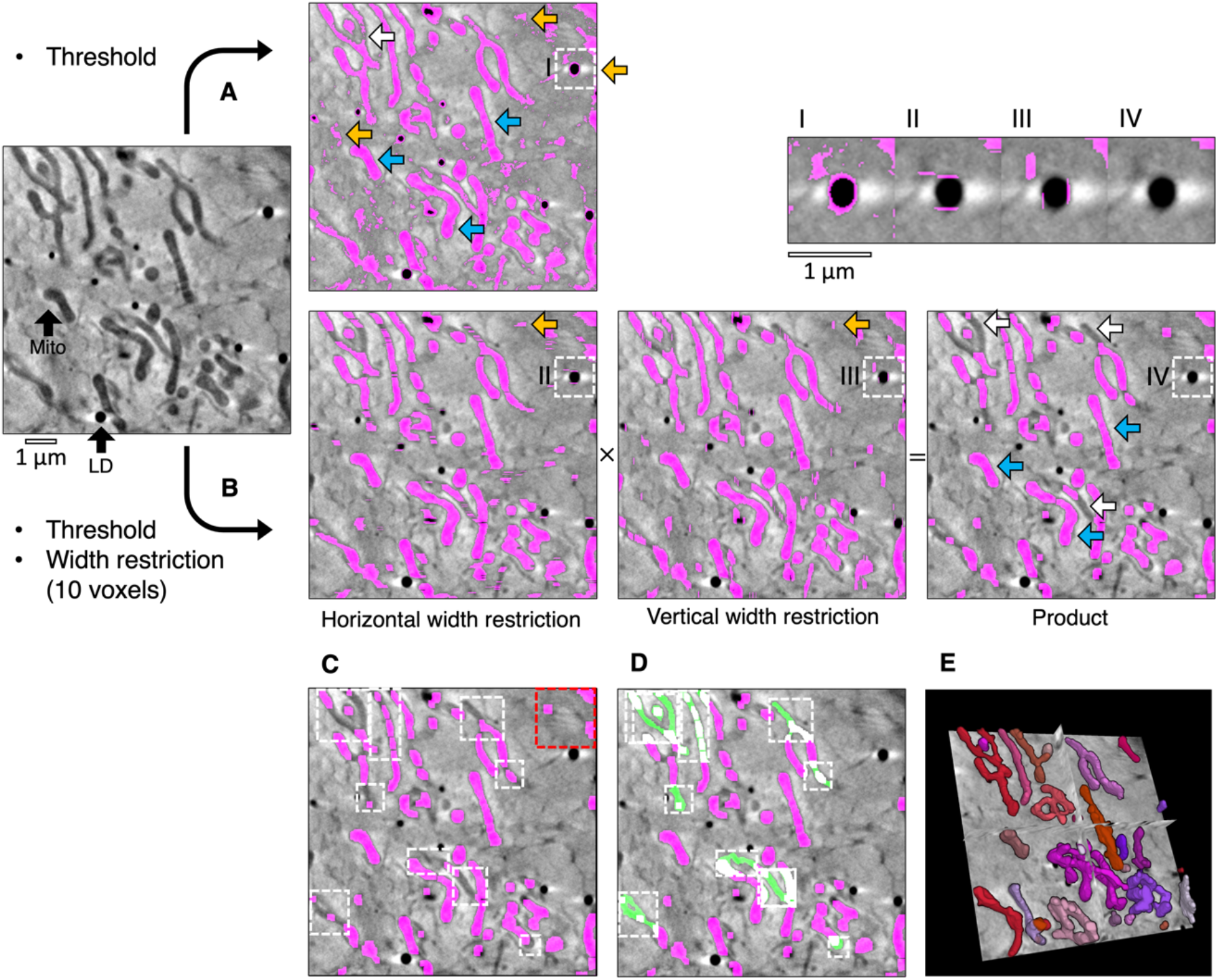
Semi-automated segmentation by analysing the intensity and width of cellular features. (A) The mitochondria in a tomogram of a U2OS cell were segmented by applying a voxel intensity threshold (blue arrows) LD, lipid droplet; Mito, mitochondrion. This technique was highly sensitive as most of the mitochondria were included and only a few areas were missing (white arrows). However, intensity thresholding alone led to noise and non-specific features such as the outline of lipid droplets being included in the segmented volume (orange arrows). (B) In *Contour*, a width restriction was applied in addition to an intensity threshold to segment the mitochondria. Any voxels included in the threshold range would only be included in the product segmented volume if they formed part of a 10×10 voxel area or larger (I-IV). The segmented product was specific to mitochondria, with less noise and fewer unwanted elements. However, there were more falsely-excluded areas due to the higher specificity (white arrows). (C) Remaining non-specific elements were manually erased (red box) and local regions of interest containing the excluded areas were identified (white boxes) and (D) the analysis was reattempted with a smaller width restriction of 4 voxels (green fill).(E) The final segmented volume was rendered in 3D using 3D Viewer in Fiji(2).

During global segmentation, the same threshold range applied in Figure 1A was applied to the tomogram in Figure 1B to isolate voxels of the desired intensity and to produce binary masks for each Z image (0 for background voxels and 1 for segmented voxels). A width restriction was determined by manually inspecting the width of the mitochondria and was applied in the second step to exclude noise and non-specific elements smaller in width than the mitochondria, such as the outline of lipid droplets. In order to apply this restriction without the slow process of iterating through each voxel, the binary masks were compressed in a lossless manner by run-length encoding^(27)^. Using this compression method, the run of voxel values (e.g. 000110000) in the binary mask were compressed into a sequence where the voxel value was coupled to the number of times it appeared consecutively (e.g. (0,3),(1,2),(0,4)). The width restriction was applied to the compressed sequence by converting voxels with a value of 1 to 0 if the number of consecutive voxels was lower than the desired width. The data compression and width restriction were applied twice independently along rows and columns in the horizontal and vertical directions, respectively, and the modified sequences were decompressed into two full binary masks. Voxels segmented within the threshold range were converted into background if their width was less than the width restriction. As a result, the segmented voxels that remained appeared as stripes with a width greater than or equal to the width restriction. The stripes were horizontal or vertical depending on the direction in which the width restriction was applied (Fig. 1B). The arrays of voxels that made up the horizontal and vertical binary masks were multiplied together such that only coordinates that contained a voxel of 1 in both masks (i.e. 1×1) were included in the product segmented volume and all other combinations were converted to background (i.e. 1×0, 0×1, and 0×0). This multiplication step eliminated most noise by ensuring that only rectangular matrices of dimensions width×width or larger remained. In some cases, horizontal and vertical stripes were produced from noise or non-specific features, such as the outline of lipid droplets. Voxels at the intersection between these stripes (i.e. 1×1) were also included after the multiplication step. The run-length encoding, width restriction, and data decompression were reapplied to the product segmented array to filter out these artefacts. The combined application of thresholding and a width restriction results in a better-defined segmentation with less noise and fewer non-specific elements. However, the increase in specificity afforded by width analysis can lead to some desired elements becoming excluded from the segmented volume. In the presented example, the global segmentation step excluded several areas based on the minimum width restriction (Fig. 1C). These areas could be filled by using the local segmentation algorithm in *Contour*, whereby thresholding and width restriction were applied locally in a smaller 3D region of interest containing these excluded areas (Fig. 1D) using a lower minimal width value (4 voxels). Given that local segmentation is performed on a smaller 3D region of interest, there is no requirement for data compression by run-length encoding before applying width restriction to improve analysis efficiency^(27)^.

It is likely that local segmentation will be required following global segmentation. However, global segmentation of the complete Z stack is not required before performing local segmentations. If it is determined that the cytoplasm is too dense with high-contrast compartments to perform a global segmentation, this step can be skipped and local segmentations can be performed on the entire tomogram instead (Fig. 2A and Table 1). In addition to the global and local segmentation algorithms, manual ‘fill’ and ‘erase’ options are available for manual adjustment of the segmented volumes (Fig. 1 C and D). The segmented volume can be rendered using 3D Viewer in Fiji^(2)^ or other appropriate visualisation software (e.g. Amira (Thermo Scientific) or Chimera/ChimeraX (UCSF)^(28)^) (Fig. 1E).

**Figure 2.**
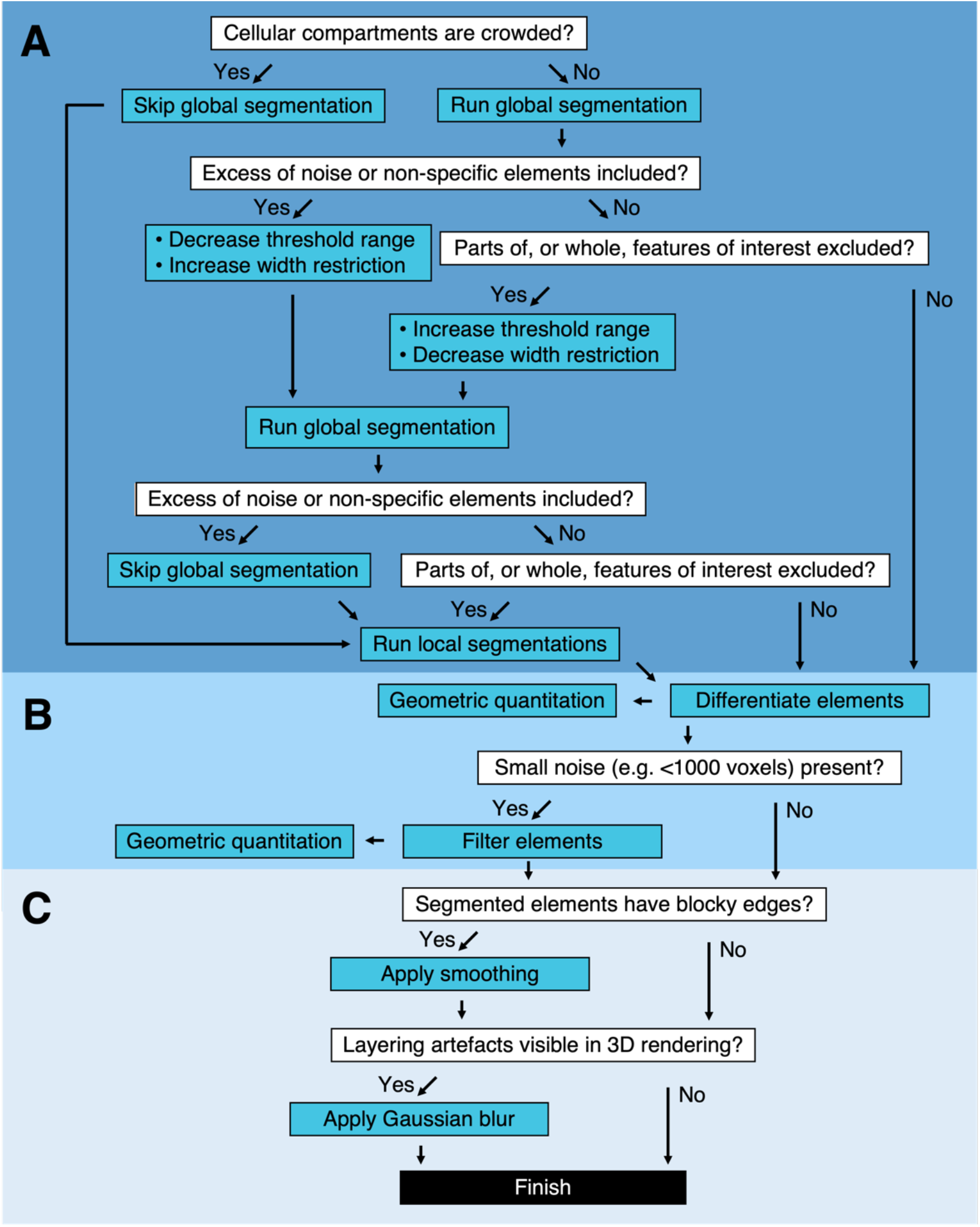
Segmentation pipeline and decision tree in *Contour*. (A) Global and local segmentation algorithms can be applied to delineate cellular compartments from a cryoSXT Z stack or from smaller 3D regions of interest. Global segmentation is recommended if the cellular compartments are dispersed throughout the tomogram. For smaller regions of interest, the local algorithm can be used to discriminate features in crowded areas or features excluded from the global segmentation. The threshold range and width restriction parameters can be modified to optimise the specificity and sensitivity of the global segmentation. (B) Discrete segmented elements can be differentiated and their volumes and widths can be calculated. Any elements smaller in volume than a specified number of voxels can be filtered out and this can be used to eliminate small segments of noise in one step. (C) Final touches can be applied to improve the appearance of the segmented volumes. A smoothing function can be used to smoothen blocky edges in 2D slices and a Gaussian blur can be applied to reduce the appearance of layering in between slices of the segmented volume (Fig. 4).

**Table 1.** Troubleshooting segmentation in *Contour*.

| Problem | Possible cause | Possible solution |
| --- | --- | --- |
| **Feature detection** |  |  |
| Global segmentation failed to produce a segmented volume | Thresholding and width restriction parameters were too stringent | Increase the threshold range and/or reduce the width restriction |
| Global segmentation produced a lot of noise | Thresholding and width restriction parameters were too permissive | Reduce the threshold range and/or increase the width restriction |
|  | Cellular compartments in the field of view were too crowded | Skip global segmentation and perform local segmentations instead |
| Multiple features of interest were excluded after the global segmentation | Thresholding and width restriction parameters were too stringent | Fill in the excluded regions using local segmentations |
|  | Cellular compartments of interest have uneven projection intensity |  |
| The edge or terminus of a feature or a constricted region within the feature was excluded from a global or local segmentation | The width restriction was too stringent at this region | Apply a local segmentation to this region with a reduced width restriction |
| **Noise elimination** |  |  |
| Too many small regions of noise (e.g. <1000 voxels) are present in the segmented volume. | Width restriction parameters were too permissive | Noise can be eliminated altogether in one step using the filter function that eliminates segmented elements below a certain volume of voxels. The elements need to be differentiated as a prerequisite. |
| **Appearance of segmented volume** |  |  |
| The segmented elements have blocky edges | A high minimum width restriction led to large width×width areas being produced in the segmented volume | Apply the smoothing function to the segmented volume |
| The segmented elements are too thin in the smoothened segmented volume | Too many iterations of the smoothing function were applied, resulting in overtrimming of the edges. | Use fewer iterations (1 to 3 are recommended) |
| Contour lines are visible in a 3D render of the segmented volume | The segmented volume was not smoothened or blurred. | Apply the smoothing function to the segmented volume and apply a Gaussian blur. |

### Applications of *Contour* to analyse geometry of cellular compartments

We have shown that mitochondria can be segmented using the global and local segmentation parameters based on their intensity and width (Fig. 1, 2, and 3A). We have used *Contour* to segment mitochondria in a recent preprint where we studied how mitochondrial morphology changes during HSV-1 infection. We found that mitochondria transitioned from a heterogenous morphology in uninfected U2OS cells to a more consistently elongated and branched formation as the infection progressed^(24)^. *Contour* can be used to segment other cellular compartments based on intensity and width, such as lipid droplets (Fig. 3B) and features at the cell surface or at cell-cell junctions, such as large internalisations of the plasma membrane that may resemble bulk endosomes arising from clathrin-independent endocytic events (Fig. 3C)^(29)^.

**Figure 3.**
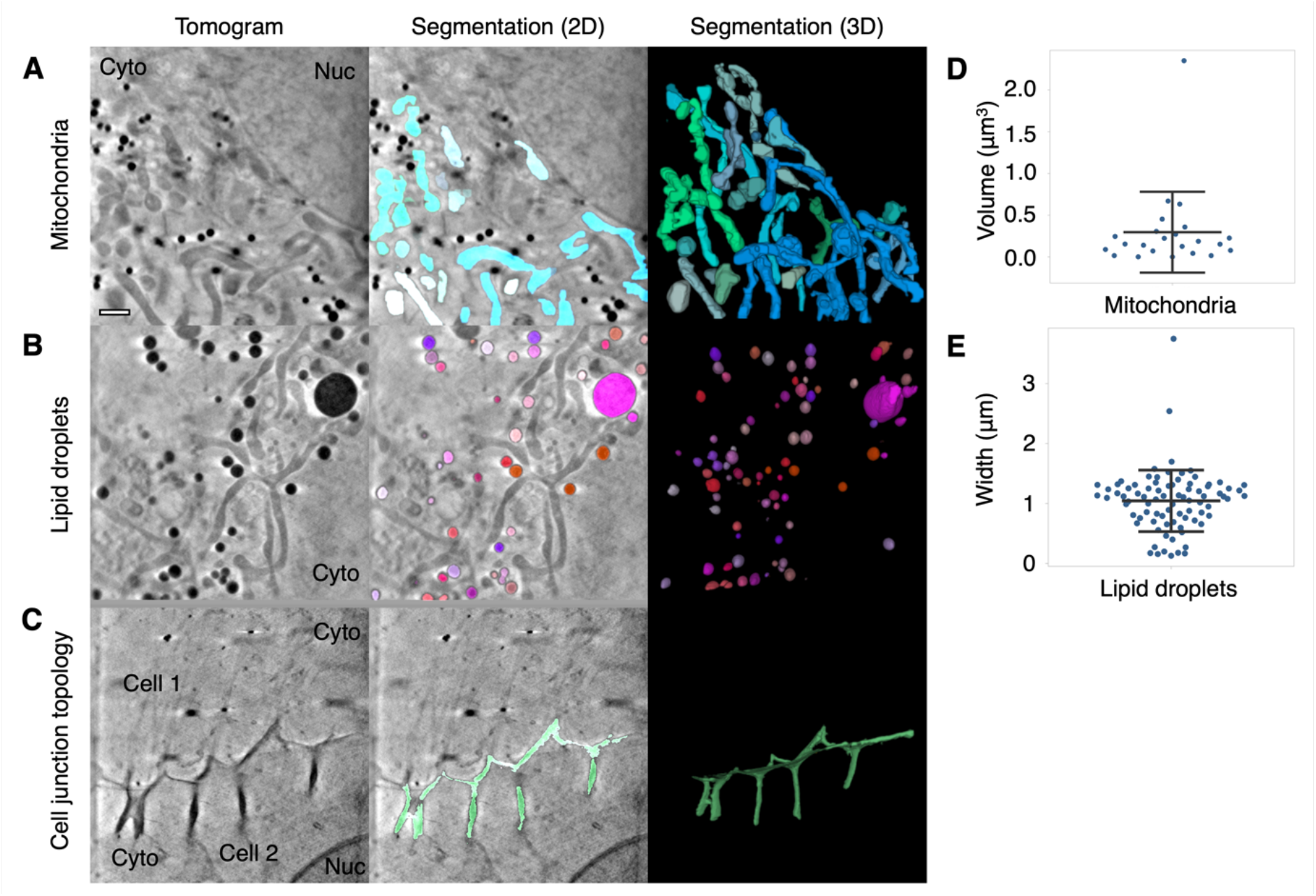
Segmentation and quantitation of cellular features. *Contour* can be used to segment high contrast features in U2OS cells such as (A) mitochondria, (B) lipid droplets, and (C) distinctive membrane topology at cell-cell junctions. Cyto, cytoplasm; Nuc, nucleus. Quantitative data can be extracted from the segmented volumes. (D) The mitochondria in this 9.46×9.46 μm^2^ field of view of a U2OS cell had a mean volume of 0.3 ± 0.48 μm^3^ SD. (E) The mean width along the longest axis of each lipid droplet in this 9.46×9.46 μm field of view of a U2OS cell was found to be 1.04 ± 0.51 μm SD. Scale bar = 1μm. Error bars show mean ± SD.

Discrete segmented elements can be differentiated from each other and colour-coded to aid discrimination of the components (Fig.2B and Fig. 4). This is achieved by assigning a common ID number to segmented voxels and their direct-contact neighbours. The inclusion criteria for direct-contact neighbours are any two voxels that are at XY coordinates that differ by one step in any of the eight cardinal (N,S,E, or W) and ordinal (NE, SE, SW, or NW) directions; or any two voxels at the same XY coordinate in tandem Z planes.

**Figure 4.**
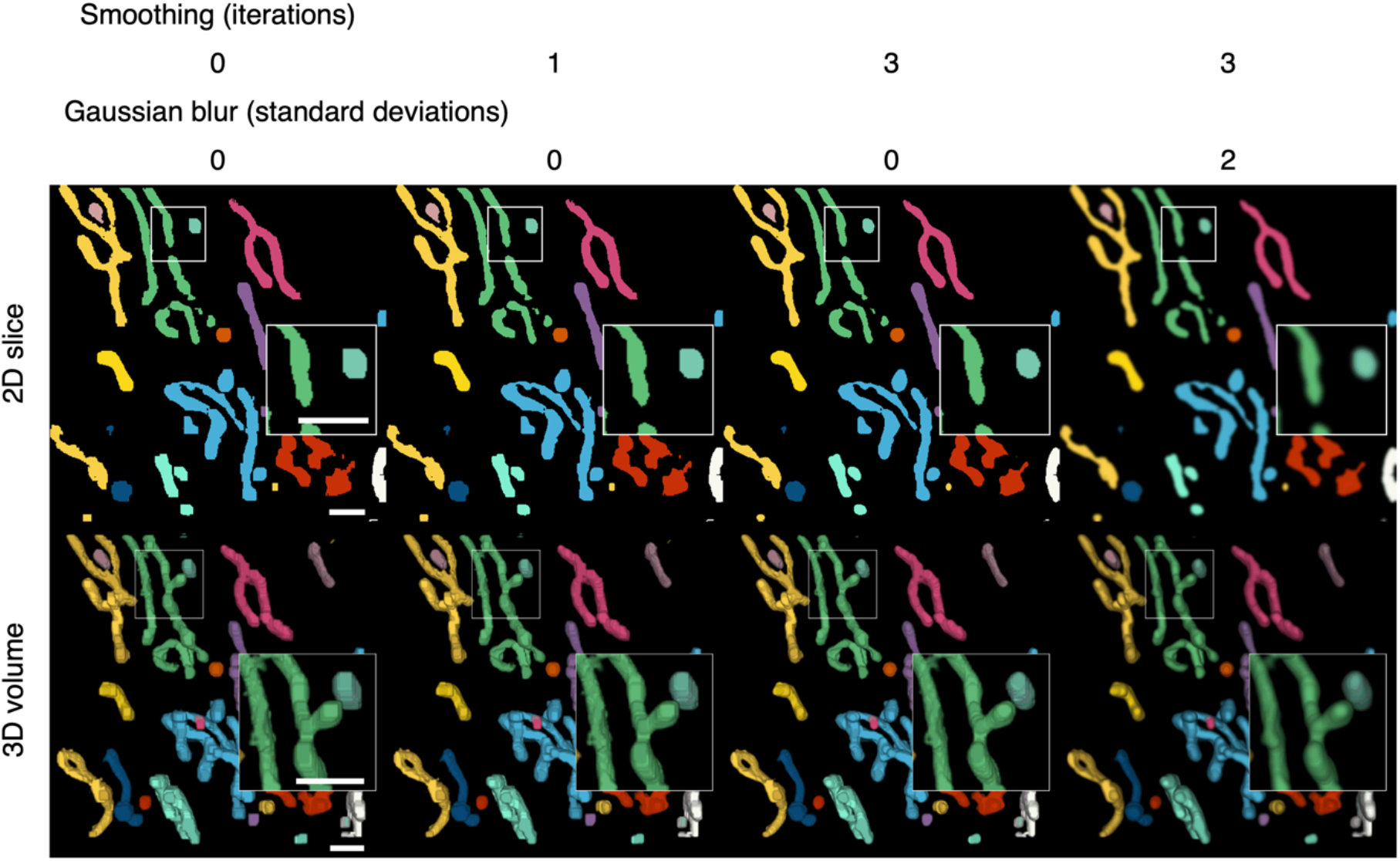
Colour-coding of differentiated elements and smoothing of the 3D volume. Segmented voxels are grouped together into separate elements that can be colour-coded to help distinguish them from each other. A smoothing function can be applied to 2D arrays of voxels to smooth the edges of segmented elements. Because the smoothing is applied to the 2D slices, layering artefacts can be observed in between the slices. A Gaussian blur can be applied per 2D slice to reduce the appearance of layering artefacts. Scale bars = 1μm.

Quantitation of the geometry of cellular features is a current challenge in cryoSXT because segmentation is often a prerequisite and measurements may need to be taken at an angle distinct from the slices of the 3D projection^(16)^. *Contour* has the capacity to automatically calculate the volumes of cellular features (in units of voxels) along any axis once the user has differentiated these elements. For example, the mean volume of the mitochondria in a single 9.46×9.46 μm^2^ field of view of a U2OS cell, given a voxel size of 10 nm^3^, was calculated to be 0.3 ± 0.48 μm^3^ (mean ± SD; Fig. 3D). The width of each segmented element along its longest axis, which may not be parallel with the slices of the tomographic projection, can also be calculated in this program. This is achieved by isolating the voxels at the perimeter of each segmented element in each image plane and calculating all combinations of the distance (i.e. modulus) between any two of these voxels across the complete Z stack. The longest of these moduli is presented as the width of the segmented element in units of voxels. The longest width of each lipid droplet was calculated for a 9.46×9.46 μm^2^ field of view and the droplet width was found to be 1.04 ± 0.51 μm (mean ± SD; Fig. 3E). Segmented volumes generated with other segmentation tools, such as Segmentation Editor in Fiji^(2)^, can be imported into *Contour* for quantitation based on the methods described above.

### Polishing the segmented volume

After the segmented elements have been differentiated, final touches can be applied to improve the appearance of the 3D volume (Fig. 4). The width restriction applied during the segmentation filters out any segmented voxels that do not form part of a width×width area or larger. As a result, segmented elements may appear blocky. A smoothing function is supplied to smoothen the edges of segmented elements (Fig. 4). Each segmented plane in the Z stack is converted into a binary mask (0 for background and 1 for segment) and is translated by one step in all eight cardinal and ordinal directions and the voxel arrays are added together such that voxels may have a value of 0 to 8. Voxels with less than a median of 5, which occur at the perimeter of segmented elements, were transformed into background, resulting in the trimming of the edges of the segmented elements. A greater number of iterations of this function increase the extent of smoothing but reduce the width of the segmented elements. A compromise of 1-3 iterations is recommended to avoid overtrimming (Table 1). The smoothing function is only applied within slices of the segmented volume and layering artefacts can be observed in between slices. A two-dimensional Gaussian blur can also be applied per slice to reduce the appearance of layering artefacts and improve the 3D rendering of the volume.

## Discussion

Here we reported the development of *Contour*, a segmentation tool for highly contrasting cellular features in cryoSXT tomograms that analyses the projection intensity (i.e. darkness) and width of cellular compartments. This program also calculates 3D geometric measurements from the segmented elements. We demonstrate that mitochondria, lipid droplets, and the topology of the cell surface at cell-cell junctions can be segmented using this technique. *Contour* was developed to accelerate segmentations of cryoSXT tomograms. Existing segmentation techniques may be time-consuming and laborious to users: manual segmentation tools require the user to trace the edges of features in periodic Z planes and interpolate between them and, although machine-learning tools such as SuRVoS are available, these tools require fresh training for each tomogram^(3,4,16)^. The algorithm used by *Contour* for segmentation is largely automated, allowing users to perform either a global segmentation on a complete cryoSXT Z stack or local segmentations in regions of interest. In either case, training datasets are not required, and the user does not need to trace around features, making the process less laborious and subjective^(4)^.

We have applied *Contour* to one study, where we investigated how HSV-1 infection alters the morphology of cellular compartments, and we were able to segment mitochondria in multiple tomograms^(24)^. The dependency on low projection intensity and width for the segmentation does pose some limitations. For example, some cellular compartments such as mitochondria may have uneven intensities. It is still possible to use *Contour* for these features, but successful analysis requires a greater number of local segmentations to be carried out with different threshold ranges (Table 1). The use of a width restriction parameter to distinguish features from noise complicates the application of this technique to thin cellular features, such as cytoskeletal filaments that are normally less than five voxels in width^(30)^. Cytoplasmic vesicles often have a highly contrasting membrane but a light lumen, making it difficult to segment such features when applying a minimum width restriction. Although we did not use *Contour* to segment cytoplasmic vesicles in our recent study^(24)^, we used *Contour* to calculate the longest widths of each vesicle that we manually segmented using Segmentation Editor in Fiji^(2)^. We therefore show that *Contour* can be used in conjunction with other segmentation tools to calculate quantitative data. Our semi-automated segmentation tool could be used to generate sufficient segmented volumes of different cellular compartments to facilitate training of machine learning algorithms in the future. CryoSXT is a growing technique and its applications are becoming more widespread in biomedical imaging, especially as a correlative imaging tool with cryoSIM^(17–19)^. *Contour* is a largely automated segmentation tool designed to keep up with the pace of tomogram acquisition and to provide a new method for quantifying tomographic data.

## Materials & Methods

### Sample preparation

3 mm gold EM grids with a holey carbon film (R 2/2, 200 mesh; Quantifoil Cat no. AU G200F1 finder, batches Q45352 & Q45353) were glow discharged and treated with filtered poly-L-lysine for 10 minutes (Sigma Aldrich Cat no. P4832). U2OS cells (ATCC HTB-96; RRID CVCL_0042) were seeded onto the grids at 3 × 10^5^ cells per well in a 6-well plate. The cells were cultivated overnight in Dulbecco’s Modified Eagle’s Medium (DMEM; Thermo Fisher Scientific, Cat no. 45011590366) supplemented with 10% (v/v) fetal bovine serum (FBS; Capricorn, Cat no. FBS-11A), 4 mM L-glutamine (Thermo Fisher Scientific, Cat# 25030081), and penicillin/streptomycin (10000 U/ml; Thermo Fisher Scientific, Cat# 15070063). 2 µL of gold fiducials (BBI Solutions; EM.GC250, batch 026935) were added to the grids as previously described^(18)^ and the grids were blotted with for 0.5-1 s at 30°C and 80% humidity with a Leica EM GP2 plunge freezer. The grids were plunged into liquid ethane and then transferred into liquid nitrogen. The tomograms presented in this paper were collected for a study of the effect of HSV-1 infection on the morphology of cellular compartments in U2OS cells^(24)^. All tomograms shown here were collected from uninfected cells except for Fig. 3B, which was collected from a cell infected with 1 plaque forming unit per cell of HSV-1 as previously described^(24)^.

### Cryo-soft-X-ray tomography

CryoSXT data was collected at beamline B24 at the UK synchrotron Diamond Light Source using a UltraXRM-S/L220c X-ray microscope (Carl Zeiss X-ray Microscopy). Soft X-rays (500 eV, λ=2.48 nm) were focussed onto the grid sample by diffraction using a diffraction grating known as a zone plate, which can achieve a nominal resolution of 25nm. A 1024B Pixis CCD camera (Princeton instruments) was used to collect tomographic data from U2OS cells with a 9.46×9.46 μm field of view by rotating the grid within the range -60° to +60° at increments of 0.5° or 1.0° and X-ray exposure times of 0.5 s or 1.0 s. A single-axis alignment of the tomographic images were generated using IMOD (version 4.9.2)^(25)^. A coarse alignment with a high-frequency cut off radius of 0.1 and a subsequent fine alignment with fiducial tracking were used to align the images. The data was reoriented in 3D using a boundary model. A final alignment was carried out using linear interpolation and tomograms were reconstructed using the back projection strategy with radial filtering to reduce noise in the form of 20 iterations of simultaneous iterations reconstruction technique (SIRT)-like filter^(26)^. The tomograms were converted from a 16-bit signed format to an 8-bit format before segmentation.

### Global segmentation

Tomographic images are stored as NumPy^(31)^ arrays in Python 3.7 and the images in the Z plane are stored in a list. Datasets with a field of view greater than 512×512 voxels were downscaled by a multiple of two to improve the efficiency of the program and the scaling was accounted for during quantitation. A threshold range with a desired minimum and maximum value was applied to produce binary masks for each image (0 for background and 1 for segmented voxels). The sequence of 0s and 1s is compressed losslessly by run-length encoding into a paired sequence where the value is coupled to the number of times it is repeated^(27)^. Values of 1 are converted to 0 if the number of repetitions is lower than the width restriction and the processed sequence is decompressed into a full array. The run-length encoding and width restrictions are applied twice independently—down columns and along rows. Both binary arrays are multiplied together so that only voxels with a value of 1 in both arrays are included in the product array. This process is repeated to remove artefacts.

### Local segmentation

A cuboidal region of interest is selected from the Z stack and a threshold range is applied to produce binary masks for each image. A width restriction is applied by iterating through the voxels in each image plane in the region of interest and counting the number of repeats. If the number of repeats is lower than the width restriction the values are converted from 1 to 0 This process is run along rows to produce a new array. This new array is used as the input array to rerun the width restriction down columns. This process is repeated once along rows and columns to remove artefacts.

### Quantitation and filtering

Segmented voxels were attributed with integer IDs that served to distinguish discrete elements. IDs were shared between neighbouring voxels that were one position away from each other in all cardinal (N,S,E, or W) and ordinal (NE, SE, SW, or NW) directions or voxels with matching XY coordinates in tandem Z planes. Neighbouring voxels were first grouped together in two dimensions in the XY planes. Any two-dimensional groups from tandem Z planes were merged into one 3D group if they contained voxels whose coordinates overlapped in XY. This process was run in ascending and descending order of Z slices to ensure that segment branches, which were separated from the main body of the segment in some slices, were not excluded from the 3D merger owing to the direction of iteration through the Z stack. The volume of each 3D segment was calculated in units of voxels. This was done by isolating all the voxels in the segmented volume with a given ID using the NumPy.argwhere function^(31)^, which produces an array of XY coordinates corresponding to these voxels per Z slice. The length of the arrays for each slice were calculated and divided by two to retrieve the number of voxels. Small segmented elements of noise were eliminated by replacing any elements with a volume of less than a desired volume threshold (e.g. 1000 voxels) with background voxels of value 0.

The width of a 3D segmented element was calculated by finding the longest distance between any two voxels in the segment. First, the voxels present at the perimeter of elements were filtered from all the voxels in the segment by determining if any neighbouring voxels have a value of 0 (background). Second, the modulus between all combinations of two perimeter voxels was calculated (Equation 1). The longest modulus was given as the width of the segment in units of voxels. Stacks of binary masks containing elements with known volumes and widths were generated to verify the quantitation functions and are available at https://github.com/kamallouisnahas/Contour/tree/main/known_quantities.

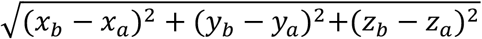

Equation 1. The modulus of all vectors connecting perimeter voxels a and b was calculated from coordinates x, y, and z.

### Smoothing and Gaussian blur

The edges of segmented elements were smoothed by translating the arrays of voxels for each slice in the tomographic projection by one voxel in each cardinal and ordinal direction. Binary masks were used and each segmented voxel had a value of 1. A sum array was produced by adding together all eight translated arrays, such that voxels ranged from 0 to 8. A median array was calculated from the sum array by transforming voxels < 5 into values of 0 and voxels ≥ 5 into values of 1. Several iterations of this function (up to 3) were applied to increase the extent of smoothing.

The Gaussian filter function from the SciPy library^(32)^ was applied with a standard deviation of 1 to each of the three colours individually in RGB images of the differentiated segmented elements. Quantitation of volume and width were not affected by the smoothing and Gaussian blur functions.

### Statistics

SuperPlots was used to generate scatterplots and to calculate the mean and standard deviation for the volume of mitochondria and the width of lipid droplets^(33)^.

## Acknowledgements

We thank Diamond Light Source for access to beamline B24 (mx18925, mx19958, bi21485 and bi23508) and the experimental hall coordinators for helpful support. We thank members of beamline B24 at the Diamond Light Source (Mohamed Koronfel, Ilias Kounatidis, Chidinma Okalo, and Matt Spink) for technical support with cryoSXT. We thank Thomas Fish (Diamond Light Source) for his assistance with NumPy array functions.

## Author Contributions

KLN, JFF, MH, CMC, and SCG conceptualized the study; KLN curated the data and visualisation; MH, CMC, and SCG provided funding; Investigation was performed by KLN, CMC, MH, and SCG; MH, CMC, and SCG administered the project; MH provided resources; KLN provided software; MH, CMC, and SCG supervised the project; KLN and MH wrote the original draft and KLN, JFF, MH, CMC, and SCG reviewed and edited the draft.

## Data Availability Statement

The raw data tilt series and tomographic reconstructions of the cryoSXT datasets presented here can be accessed from the Apollo repository (University of Cambridge): https://doi.org/10.17863/CAM.78593. The source code is available under a GNU General Public License v3.0 from https://github.com/kamallouisnahas/Contour. The segmented volumes and quantitative data can be accessed from: https://github.com/kamallouisnahas/Contour/tree/main/repository.

